# Effect of Spatial Release from Masking on Listening Effort in Different Semantic Contexts

**DOI:** 10.64898/2026.07.13.738303

**Authors:** Agudemu Borjigin, Nimesha Didulani Dantanarayana, Yuhang Li, Ruth Y. Litovsky

**Affiliations:** University of Wisconsin-Madison (Madison, WI, USA); University of Utah (Salt Lake City, UT, USA)

**Keywords:** listening effort, spatial release from masking, semantic context

## Abstract

Humans often communicate and learn in noisy, complex listening environments. Here, we investigated the effects of spatial hearing and semantic context cues on speech intelligibility and listening effort in young adults with typical hearing. The listening task included conditions in which target speech and speech maskers were either spatially co-located or separated. Target sentences were either semantically coherent or anomalous, while the masker comprised a mixture of two coherent sentences. Results showed higher speech intelligibility in spatially separated than co-located conditions, demonstrating a robust spatial release from masking (SRM), which is consistent with prior findings. SRM did not differ between semantically coherent and anomalous sentences, indicating comparable benefits of spatial cues across semantic contexts. However, within each spatial configuration, intelligibility was higher for coherent than anomalous sentences. Listening effort, indexed by peak pupil dilation in pupillometry measurement, was reduced in spatially separated conditions, suggesting a trend toward a release from listening effort. Analysis of the timing of peak pupil dilation revealed a significantly delayed peak dilation for anomalous sentences in the co-located condition compared with coherent sentences in the separated condition, indicating increased processing demands in the absence of spatial and semantic cues. Finally, SRM was correlated with the magnitude of release from listening effort for coherent sentences, but not for anomalous sentences, suggesting that intelligibility and listening effort benefits might co-occur when contextual cues are available.

## 1 Introduction

The ability to selectively attend to target speech while ignoring interfering talkers is a complex skill used in everyday listening situations and represents a feature of the “Cocktail Party Effect” (Cherry, 1953; Pollack & Pickett, 1958). When listening to speech in noise, spatial cues play an important role in significantly improving speech recognition. An improvement in speech recognition is typically observed when the target speech signal and masking noise are spatially separated compared to spatially co-located, which is commonly termed spatial release from masking (SRM) and shown by many studies (Drullman & Bronkhorst, 2000; Hawley et al., 1999; Plomp & Mimpen, 1981; Srinivasan et al., 2024; Zenke & Rosen, 2022). In addition to spatial cues, listeners also use contextual cues to enhance speech recognition by predicting words they missed due to speech masking (Tavano & Scharinger, 2015). Language predictions incorporate low-level acoustic details and high-level processing of language content. Top-down high-level processing is especially important in complex listening situations where bottom-up sensory processing is compromised due to noise masking (Rönnberg et al., 2013). Top-down compensation interacts with bottom-up processing of acoustic–phonetic features of the speech, and both processes also interact with the listener’s long-term linguistic and semantic knowledge during speech recognition in complex listening situations (Luce & Pisoni, 1998; Tuennerhoff & Noppeney, 2016). Therefore, the segregation of target speech from background noise depends not only on peripheral auditory processing, but also on higher-level cognitive processing (Rönnberg et al., 2013). In this study, we examined the contribution of higher-level cognitive processing to speech recognition in noise and its relationship with SRM by manipulating both the semantic context of the speech material and target-masker spatial configurations. We used semantically coherent versus anomalous sentences to manipulate the semantic context (Davis et al., 2011). Semantically coherent sentences allow listeners to use contextual cues of speech with top-down processing, while semantically anomalous sentences block the use of such cues (Pisoni & Kronenberger, 2021).

We also evaluated speech recognition beyond speech intelligibility scores by measuring pupil dilation as a proxy for listening effort, which reflects cognitive processing in listening-related tasks (Abdel-Latif et al., 2025; Winn et al., 2018; Winn & Teece, 2021). Speech intelligibility scores alone do not reflect individual differences in higher-up processing: two individuals with the same percent correct scores could have exerted different amounts of listening effort (Koelewijn, Zekveld, Festen, & Kramer, 2012; Mackersie & Cones, 2011). The evaluation of listening effort has the potential to add additional insight beyond speech recognition scores alone. According to the Framework for Understanding Effortful Listening (Pichora-Fuller et al., 2016), listening effort is defined as the deliberate allocation of available resources when carrying out a listening task to overcome obstacles that make task completion more difficult in goal pursuit. In recent years, there has been an increasing number of studies evaluating listening effort in addition to the percent correct scores (Ohlenforst et al., 2017; Peelle, 2018; Strauss & Francis, 2017). Listening effort has been studied using both behavioral and physiological measures (Ohlenforst et al., 2017). Behavioral assessments include subjective rating scales (Gatehouse & Noble, 2004; Rudner et al., 2012), reaction time (Voola et al., 2024), and working memory (DiGiovanni et al., 2017). Although proven effective, these behavioral approaches may not be sufficiently sensitive to reflect individual differences in listening effort exertion (Zekveld et al., 2018).

In this study, pupillometry was used to assess listening effort. Task-related changes in pupil dilation are thought to provide a direct and reliable measure of cognitive load and arousal linked to the locus coeruleus-norepinephrine system that modulates attention (Beatty, 1982; Gabay et al., 2011; Just et al., 2003; Koelewijn, Zekveld, Festen, & Kramer, 2012; Van Der Meer et al., 2010). Pupillometry is a time-series measurement rather than only capturing an endpoint measurement taken at the end of the stimulus, such as reaction time measurements and categorical rating scales. Pupillometry could sensitively reveal the nature of incremental word-by-word perceptual events over time in sentences (Winn, 2016; Winn & Teece, 2021). The size of the pupil has been shown to increase with task demands (Engelhardt et al., 2010), intelligibility levels (Burg et al., 2021; Trau-Margalit et al., 2023; Zekveld et al., 2010), sentence complexity (Kadem et al., 2020; Piquado et al., 2010), lexical competition (Kuchinsky et al., 2013), and types of maskers (Koelewijn, Zekveld, Festen, Rönnberg, et al., 2012; Villard et al., 2023). High working memory capacity and better linguistic closure ability are associated with large pupil dilation and delayed peak pupil responses (Zekveld et al., 2011; Zekveld & Kramer, 2014), indicating an allocation of more cognitive resources associated with intensive mental processing in difficult listening situations (Koelewijn, Zekveld, Festen, Rönnberg, et al., 2012; Van Der Meer et al., 2010).

Here, we measured both speech intelligibility and pupillometry, with and without target-masker spatial separation, and with semantic context cues in the target sentences being either coherent or anomalous. The combined effects of semantic context and spatial separation cues on listening effort have not been investigated in previous studies. In Winn (2016), a rapid reduction in listening effort was observed in a group of typical hearing (TH) listeners with sentences of higher semantic context than with lower semantic context. However, this study presented sentences to listeners only in quiet settings. Zekveld & Kramer (2014) and Thakkar et al. (2025) tested speech recognition in noise and studied the relationship between listening effort (by pupillometry) and SRM, but without explicit control of the semantic context in the sentence materials. In real-world listening, the acoustic and semantic context of speech can be closely linked to support robust speech recognition in challenging listening environments and could also reduce listening effort. In this study, we hypothesized that improvement in speech intelligibility due to semantic context, along with SRM, can occur simultaneously with a reduction in listening effort, indicating shared cognitive mechanisms for semantic cue processing and listening effort along with SRM. Moreover, top-down processing may reduce listening effort and enhance bottom-up acoustic processing when semantic context is available, resulting in improved speech recognition in challenging listening environments. However, in the absence of semantic cues, we predicted that listeners may adapt by shifting to rely more on bottom-up processing of spatial cues to segregate the target speech from maskers. Based on these hypotheses, we predicted that both semantic context and spatial separation would improve speech intelligibility while reducing listening effort. Specifically, sentences with semantic context were expected to yield higher recognition scores and smaller pupil responses than sentences without semantic context. Similarly, spatially separated target and masker speech were expected to improve recognition and reduce pupil responses relative to the co-located condition. Because listeners may rely more strongly on bottom-up spatial cues when semantic cues are unavailable, we further predicted that the benefit of spatial separation would be greater for sentences without semantic context. Accordingly, peak pupil dilation was expected to be largest and latest in the co-located condition without semantic context, and smallest and earlier when both spatial and semantic cues were available.

## 2 Materials and Methods

### 2.1 Participants

We recruited 13 participants with typical hearing (TH) for this study, ranging in age from 18 - 22 years (mean = 19.8, std = 1.2). All participants are native English speakers from the USA. The test lasted about two hours for this study. Participants received a stipend or extra course credit for their time. All experimental procedures followed the regulations established by the National Institutes of Health and were approved by the Institutional Review Board (#2016-0226) of Health Sciences of the University of Wisconsin-Madison. Participants were not screened for the use of substances that could affect physiological responses measured with pupillometry.

### 2.2 Experimental Design and Statistical Analyses

#### 2.2.1 Experiment setup

Experiments were carried out in a standard sound booth (IAC Acoustics, IL, USA). Participants sat at a table with their foreheads supported on a headrest for stability during the tests. The height of the table and chair was adjusted for each participant to ensure comfort. A computer screen was placed approximately 65 cm away from the headrest on the table. The eye tracker (Eyelink 1000 Plus; SR Research, Ontario, Canada) camera was mounted to the table using an 8 cm desktop mount in front of the monitor. The illumination of the test room was set constant for all participants using two floor lamps with three-position switches in the corners of the room behind the eye tracker camera. Each lamp switch was set to provide “low to moderate” illumination. The size of the pupil was measured in pixels using the “Area” setting on the eye tracker and a sampling rate of 1,000 Hz. Stimuli were presented from a loudspeaker (Tannoy, Coatbridge, Scotland) positioned at 0- and 90-degrees azimuth.

#### 2.2.2 Stimuli and procedure

The participant’s task was to listen to recorded male-voiced target sentences in the presence of competing recorded male-voiced 2-talker mixtures. There were two types of target sentences: coherent and anomalous sentences (Davis et al., 2011). Both types of sentences were phonetically, lexically, and syntactically balanced paired sentences with equal length, consisting of 7-10 words. Coherent sentences contain contextual cues (e.g., “He ironed his shirt before he wore it”), while anomalous sentences do not (e.g., “He jilted his coast before he drew it”). For the two-talker mixture, the stimuli were drawn from the IEEE corpus (Rothauser et al., 1969). All stimuli were calibrated at 70 dB SPL. The target and masker were presented at a signal-to-noise ratio (SNR) of 1 dB. This SNR was selected based on pilot testing to provide a task difficulty level at which listeners could reliably identify the target while avoiding ceiling or floor effects in performance. The same SNR was used for both types of sentences to facilitate direct comparisons of speech recognition performances between the two types of sentences. The stimuli were presented through a loudspeaker using a high-speed audio interface (RME Fireface, Haimhausen, Germany). The duration of the sentence ranged from 4,000 to 6,000 ms. We used a custom script in MATLAB (The MathWorks, Natick, MA, USA) with PsychToolbox 3 to deliver stimuli and collect data. The target sentences were presented via the loudspeaker at 0 degrees, azimuth (in front of the listener), while the 2-talker masker was either co-located with the target at 0 degrees, or presented from a loudspeaker at +90 degrees (separated towards the right ear of the listener). Speech intelligibility was defined as the number of correctly repeated words. Pupil size was simultaneously measured using the eye tracker. Before testing, participants completed target-only practice trials to become familiar with the target male speaker’s voice and practiced the task under both test conditions (i.e., co-located and separated). The sentences used for practice were excluded from the test corpus.

During testing, participants were asked to fix their gaze on a gray cross at the center of the computer screen and to focus on the target sentences presented from the front loudspeaker while ignoring the interfering masker regardless of where it was presented. The target stream was cued by its earlier onset relative to the masker, the +1 dB SNR, and prior practice with the target male speaker’s voice. Participants’ task was to repeat the target sentence after the response cue. Before each trial, the gray cross changed its color to white to signal the beginning of the trial. This was followed by a 2,000 ms pre-trial interval, and then the trial began with a 1,000 ms baseline pupil measurement in silence before the stimulus presentation. Participants were instructed to wait for a 2,000 ms following offset of the stimulus before repeating the sentence. The end of the waiting period was indicated by the cross turning green, accompanied by two sound beeps. The target sentences were randomly selected from the corpus without replacement. The experimenter waited 10 to 15 s between trials to allow the pupil to return to baseline before initiating the start of the next trial. Participants were asked to reduce their blinking while the cross was present on the computer screen to increase the reliability of tracking pupil dilation. The testing was carried out in blocks such that each block contained 5 sentences of the same type (coherent vs. anomalous) in the same spatial configuration (co-located or separated). We adopted smaller blocks of only five sentences to avoid eye fatigue. Each sentence type and spatial configuration combination contained 6 blocks. The order of block presentation was randomized across participants. Longer breaks were scheduled after a few blocks of testing.

#### 2.2.3 Analysis of pupillometry data

Pupil data were preprocessed to eliminate artifacts and exclude noisy trials. Pupil tracks with more than 45% blinks were removed from the analysis (Burg et al., 2021). This criterion was chosen to be inclusive of data compared to more commonly used restrictive criteria (e.g. 15%, 30%) (Zekveld &Kramer, 2014; Winn et al., 2018)Previous work has shown a positive correlation between task difficulty and blink percentage: an overly strict threshold such as 15% could lead to the exclusion of a greater number of difficult trials, potentially biasing results (Burg et al., 2021). To calculate the percentage of blinks in a trial, we did not consider samples from the response period due to artifacts from the motor response (Privitera et al., 2010; Winn et al., 2015). Blinks were identified by marking samples that fell more than three standard deviations (SDs) below the mean pupil diameter (Zekveld et al., 2010). According to the best practices recommended by Winn et al. (2018), trials exhibiting irregular baselines, severe distortions, or unusually large fluctuations (growth) inconsistent with task-induced pupil dilation were also excluded.

#### 2.2.4 Statistical Analyses

Statistical analysis was performed with RStudio (R version 4.3.1). Speech intelligibility scores were transformed from percent correct to rationalized arcsine units (RAU) to mediate the ceiling effect (Studebaker, 1985). To test the prediction that both listening conditions (co-located and separated) and contextual cues (coherent sentences and anomalous sentences) affect speech intelligibility, we used a linear mixed-effects model (lme4 package, version 1.1.31) with speech intelligibility scores being the dependent factor and listening conditions and contextual cues being independent factors with random effects to account for the variability as-sociated with participants: *model* = *lmer*(*percentcorrectscores listeningconditions contextualcues*+ 1*|participant*). Similarly, we also analyzed these effects on listening effort (maximum proportional change in pupil dilation or peak pupil size during the silent period): *model* = *lmer*(*peakpupildilation listeningconditions contextualcues* + 1*|participant*). The anova function of the car package (version 3.1-2) was used to compute Type III sequential sums of squares to evaluate the predictive contributions of the independent variables and their interactions within the linear mixed-effects model. The normality of residuals was visually assessed by comparing quantile-quantile plots of the model residuals against a theoretical normal distribution, as well as statistically using Shapiro-Wilk tests. Homogeneity of the variance was evaluated by Levene’s test applied to the residuals. Post-hoc comparisons were performed using the emmeans package (version 1.8.9) to estimate marginal means, with Benjamini-Hochberg corrections applied to control the false discovery rate (Benjamini & Hochberg, 1995). Correlational analyses were performed to examine potential relationships between change in speech intelligibility (RAU), i.e., SRM, and listening effort (pupil dilation) from co-located to separated conditions. Due to our directional hypothesis that SRM leads to a release from listening effort, a one-sided test was used to evaluate this relationship.

## 3 Results

### 3.1 Speech Intelligibility Scores and Spatial Release from Masking

Figure 1 shows the absolute percent correct scores across all conditions (part a) and spatial release from masking (SRM) for two target sentence types (i.e., coherent vs. anomalous; part b)There was a significant main effect on speech intelligibility scores for both spatial configuration (*F* = 389.44*, p <* 0.0001) and sentence materials (*F* = 92.25*, p <* 0.0001); see Figure 1a. There was no significant interaction between spatial configuration and sentence materials (*F* = 0.001*, p* = 0.993). Note that the comparison of the quantiles from the model residuals with a sample normal distribution verified that the residuals follow a normal distribution. We quantified SRM as the difference in Estimated Marginal Means (EMMs) of percent-correct scores between the spatially separated and co-located conditions. SRM, computed as a difference between co-located and separated conditions, was 30.1% (SD = 7.1%) with coherent sentences. (*EMM* (*separated – co-located*) = 0.301*, p <* 0.0001). The SRM is 30.2% (SD = 8.4%) with the anomalous sentences: *EMM* (*separated – co-located*) = 0.302*, p <* 0.0001. There was no significant main effect of sentence type on SRM (*F* = 0.0002*, p* = 0.99). However, coherent sentences led to better performance than anomalous sentences: there was an increase in scores of 14.7% from anomalous to coherent sentences (*EMM (coherent - anomalous) = 0.147, p <* 0.0001) in both separated (SD = 6.8%) and co-located (SD = 7.6%) conditions.

**Figure 1.**
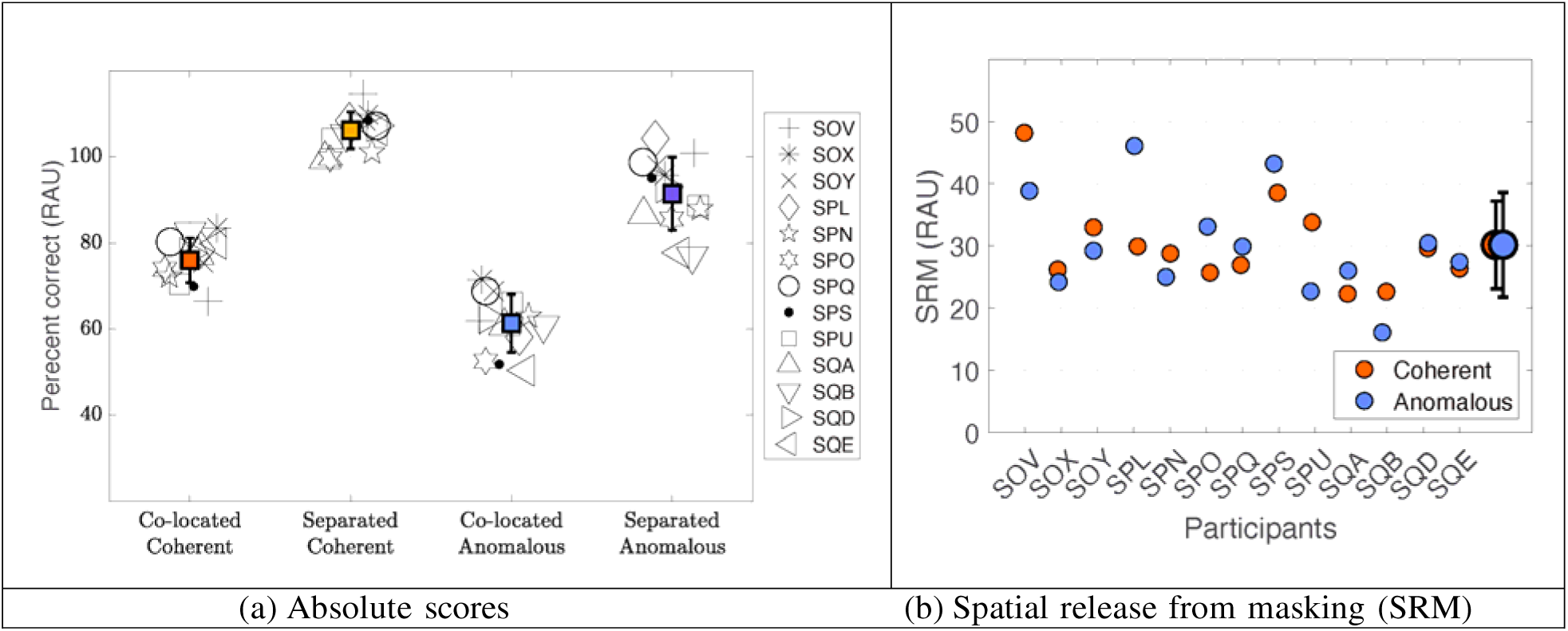
(a) Absolute speech intelligibility scores shown as percent correct scores from all four conditions. Note that warm colors (orange and yellow) represent conditions tested with coherent sentences, whereas cold colors (blue and purple) represent conditions tested with anomalous sentences. (b) SRM scores are shown for each participant.

### 3.2 Pupillometry measurements and release from listening effort

Figure 2 (top) shows grand average pupil dilation trajectory in four conditions. Peak pupil dilation and peak latency were extracted for each individual and are summarized in Figure 3. Table 1 summarizes the statistical analyses examining the effects of spatial configuration and sentence type on pupil dilation size and latency.

**Figure 2.**
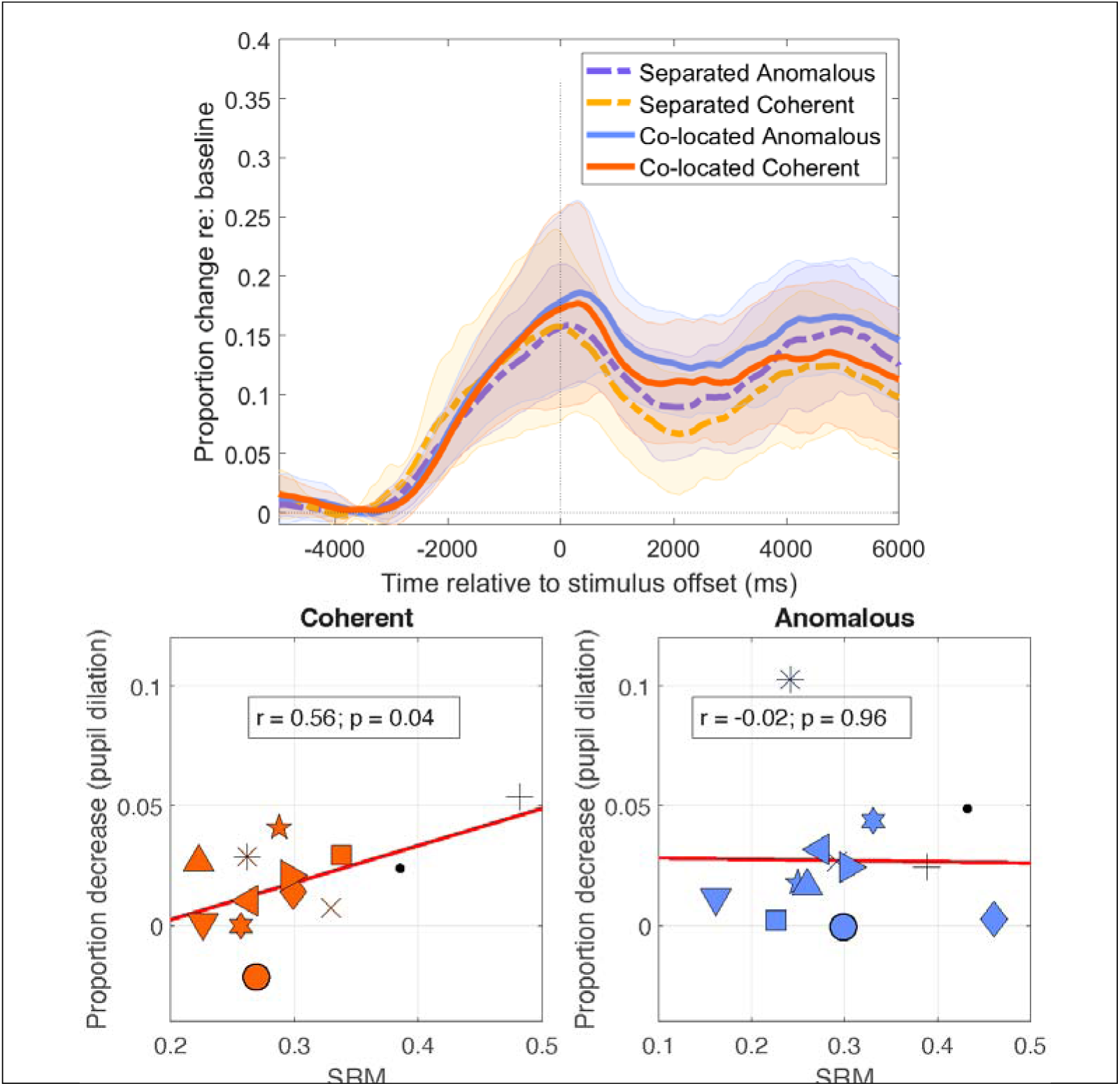
Average pupillometry data across all participants (top) and correlations between SRM and proportion decrease in peak pupil dilation due to spatial separation between target and maskers (bottom).

**Figure 3.**
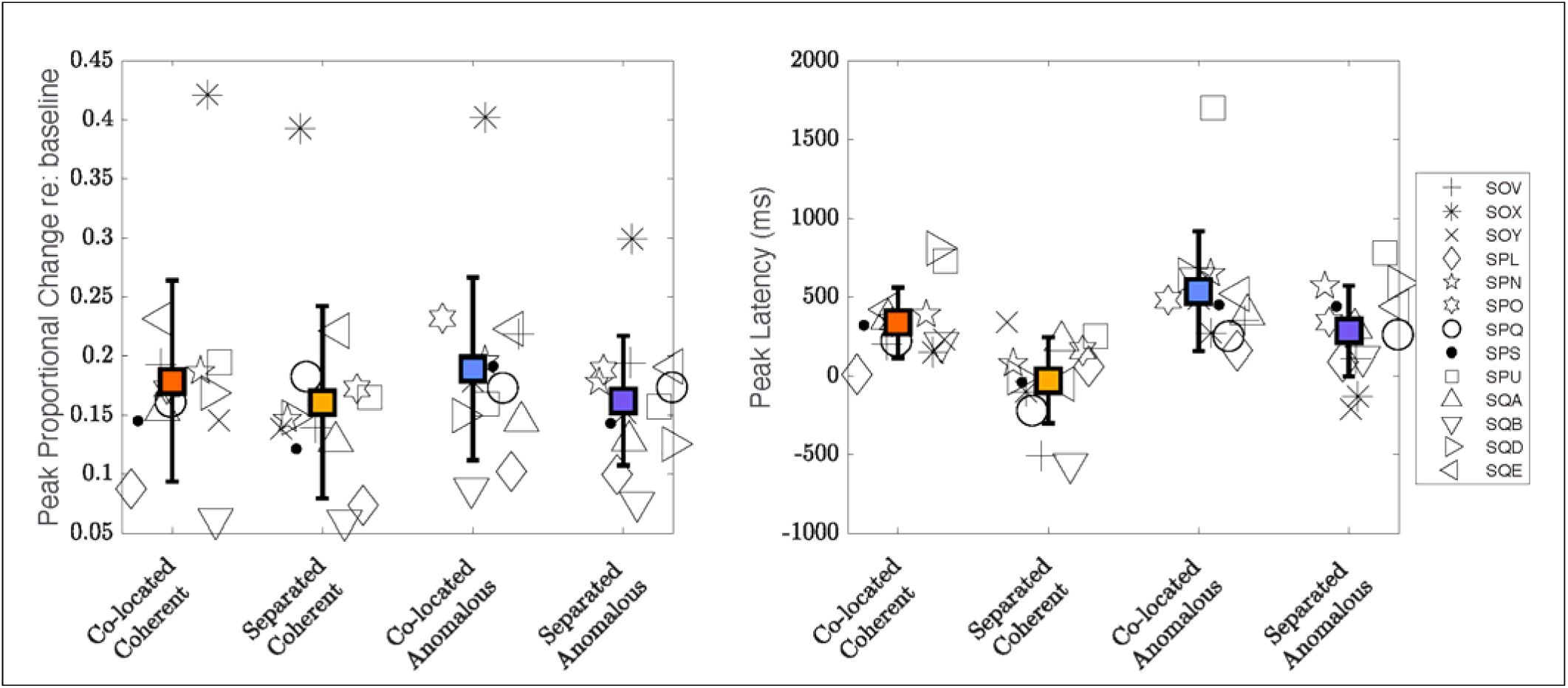
Individual peak pupil dilation (left) and peak latency (right).

**Table 1.** Estimated Marginal Means (EMM) difference in peak pupil dilation between conditions. Amplitude is in proportion change relative to baseline, while latency is in milliseconds. Statistically significant comparisons are shaded in green color.

| **Metric** | **Comparison** | **EMM diff** | **p value** |
| --- | --- | --- | --- |
| amplitude | Co-located (C) - Separated (C) | 0.018 | 0.94 |
| amplitude | Co-located (A) - Separated (A) | 0.027 | 0.83 |
| amplitude | Anomalous (Co) - Coherent (Co) | 0.01 | 0.99 |
| amplitude | Anomalous (Sep) - Coherent (Sep) | 0.0012 | 1 |
| amplitude | Co-located (A) - Separated (C) | 0.029 | 0.82 |
| amplitude | Co-located (C) - Separated (A) | 0.017 | 0.95 |
| latency | Co-located (C) - Separated (C) | 366.9 | 0.043 |
| latency | Co-located (A) - Separated (A) | 251.5 | 0.22 |
| latency | Anomalous (Co) - Coherent (Co) | 198.5 | 0.41 |
| latency | Anomalous (Sep) - Coherent (Sep) | 313.8 | 0.094 |
| latency | Co-located (A) - Separated (C) | 565.4 | 0.0011 |
| latency | Co-located (C) - Separated (A) | 53.1 | 0.97 |

Analyses of pupil dilation size did not reveal any statistically significant main effect of spatial configuration (*F* = 1.27*, p* = 0.27) or sentence materials (*F* = 0.083*, p* = 0.77). We also analyzed the peak latency, or the time point at which the peak pupil dilation occurred (relative to the stimulus offset). There was a statistically significant main effect of both spatial configuration (*F* = 15.35*, p* = 0.0003) and sentence materials (*F* = 10.53*, p* = 0.0021) on peak latency. There was no significant interaction between spatial configuration and sentence materials (F = 0.53, p = 0.47). Note that the comparison of the quantiles from the model residuals and a sample normal distribution verified that the residuals follow a normal distribution. We observed an earlier peak pupil dilation under the separated conditions than under the co-located conditions (*EMM* (*separated* −*co-located*) = − 309*, p* = 0.003). When comparing the separated vs. co-located conditions, for the anomalous sentences, pupil dilation reached its peak 251.5 ms (SD = 292.9) earlier with the separated than the co-located condition; for the coherent sentences, pupil dilation reached its peak 366.9 ms (SD = 291.4) earlier for the separated vs. co-located conditions. The peak pupil dilation also occurred earlier with coherent sentences than with anomalous sentences (*EMM* (*coherent* − *anomalous*) = − 256*, p* = 0.0094). The pupil dilation reached its peak 198.5 ms (SD = 269.1) earlier with coherent vs. anomalous sentences in the co-located conditions, and 313.8 ms (SD = 352.5) earlier in the separated conditions. There was no statistically significant correlation between SRM (i.e., speech intelligibility) and the decrease in latency due to spatial cues (from co-located to the separated condition), with either coherent sentences (*r* = − 0.18; *p* = 0.55) or anomalous sentences (*r* = 0.48; *p* = 0.1). There was no statistically significant main effect of sentence type on the decrease in latency due to spatial cues (*F* = 1.1*, p* = 0.3).

Figure 2 also shows the correlation between reductions in pupil size due to spatial cues and SRM for both coherent and anomalous target sentence types. Moderate correlation was found between SRM and the reduction of pupil dilation due to spatial cues (from the co-located to the separated condition), only with coherent sentences (*r* = 0.56; *p* = 0.04), not with anomalous sentences (*r* = − 0.02; *p* = 0.96). There was no statistically significant main effect of sentence type on pupil dilation due to spatial cues (*F* = 1.34*, p* = 0.27).

## 4 Discussion

### 4.1 Summary of Findings

This study investigated the effect of spatial hearing and semantic context cues on listening effort in young adults with TH, along with speech intelligibility measurements. As predicted, we observed higher speech intelligibility scores with spatial separation cues than without spatial separation cues, which is commonly referred to as spatial release from masking (SRM). SRM was similar for both types of sentences, indicating that spatial hearing cues improved speech intelligibility to the same extent despite differences in semantic contexts. However, within the same spatial hearing condition, we observed better speech intelligibility scores with semantically coherent sentences than with anomalous sentences. In terms of pupil dilation (used here as a proxy for listening effort), there was no statistically significant effect for spatial or semantic cues. We also analyzed the timing of when the pupil dilation reached its peak. We found that pupil dilation was significantly delayed with anomalous vs. coherent sentences, and in co-located vs. separated conditions. This latter result indicates that a longer processing time was needed in the absence of semantic or spatial cues. Lastly, we observed a moderate correlation between SRM (effect of spatial separation on speech intelligibility) and release from listening effort (effect of spatial separation on pupil dilation) with coherent sentences, but not with anomalous sentences, indicating that SRM and release from listening effort co-occur when contextual cues are available.

### 4.2 Speech Recognition and SRM in Varying Semantic Context

One of the aims of t h e current study was to examine whether and how contextual information affects listening effort and speech intelligibility in speech-on-speech masking. We sought a moderate task difficulty level so that performance was neither too low nor at ceiling across conditions. This is important for measuring listening effort because it depends on a critical combination of task demand and motivation. Listening effort diminishes if the task is too easy or too difficult, and motivation is low (Winn et al., 2018; Zekveld et al., 2018). With a male target set at 70 dB SPL and a two-talker male speech masker fixed at +1 dB SNR, speech recognition in the most difficult condition, co-located anomalous condition, was around 60%. This outcome is comparable to that reported by Abdel-Latif et al. (2025). They used semantically unpredictable low-context sentences with a single-talker male masker in a co-located setup at multiple SNRs, including 0 dB SNR with target sentences presented at 65 dB SPL. The median percentage of correctly recognized target words was around 70%. Despite small differences in target level and SNR, the similarity in performance demonstrates the robustness of the approach, ensuring a challenging but feasible task that avoids the influence of reduced motivation affecting pupil responses (Zekveld et al., 2018).

As expected, we observed significantly better speech recognition with coherent sentences than with anomalous sentences in both co-located and separated spatial configurations, indicating the importance of semantic context in speech recognition in challenging listening situations with or without spatial cues. This is in line with the study by Davis et al. (2011), which showed significantly higher speech recognition with coherent than anomalous sentences in adults with TH (tested from −7 dB to +1 dB SNRs, in increments of 1 dB). However, they tested speech recognition when the target and masker were co-located, and the masker was non-speech noise (speech-spectrum signal-correlated noise). In another study, young adults aged 22 to 28 years performed significantly better with high-context sentences than with low-context sentences in broadband noise presented at +20 dB SNR and +40 dB SNR, also only in a co-located target-masker configuration (Dubno et al., 2000). Wasiuk et al. (2022) tested speech recognition in both speech-shaped noise and a two-talker masker in the co-located condition, and observed that the semantic context provided by meaningful target sentences helped listeners more in speech-shaped noise than in the two-talker speech masker. Calandruccio et al. (2018) used very similar materials as our study: coherent and anomalous sentences as the target, and IEEE sentences as two-talker maskers. Their results also showed better performance with semantic context than without semantic context. However, similar to previous studies, Calandruccio et al. (2018) only tested the co-located condition.

In this study, we tested both co-located and separated conditions and observed a significant amount of SRM with both coherent and anomalous sentences. The current study, along with prior studies, demonstrates the effect of semantic context on speech recognition, indicating that adult listeners effectively utilize semantic context to recognize speech in difficult listening situations. This further emphasizes how listeners compensate for the loss of bottom-up information processing when recognizing speech in noise by using top-down processing associated with semantic cues (Kathleen Pichora-Fuller, 2008). In addition, the current study revealed how spatial cues are equally beneficial in the presence or absence of semantic cues, indicating that spatial cues may play a role in both top-down and bottom-up processing to facilitate speech recognition in unfavorable listening situations.

### 4.3 Listening Effort and Spatial Hearing Cues

In our study, we observed a moderate positive correlation between SRM and reduction in pupil dilation when the target speech was coherent in nature. This is consistent with the recent findings from Borjigin and Bharadwaj (2025). Their results indicate a correlation between reduced listening effort and better sensitivity to the processing of temporal fine structure of sounds, which underlies the processing of spatial hearing cues (Borjigin et al., 2022; Klumpp & Eady, 1956; Wightman & Kistler, 1992). Similar to our study, Rennies and Kidd (2018) and Rennies et al. (2019) reported a significant negative relationship between listening effort and SRM in young adults with typical hearing. In these studies, spatial release from listening effort was observed at positive SNRs where speech intelligibility was at the ceiling, showcasing the added value from measuring listening effort. However, these studies did not report spatial release from listening effort in the SNR region where intelligibility was below the ceiling. In our study, we observed a significant release from listening effort due to spatial cues when speech intelligibility was below ceiling in the spatially co-located conditions and at the ceiling in the spatially separated conditions. In addition, the studies by Rennies and colleagues used a categorical rating scale to measure listening effort, unlike the objective measure, pupil dilation, used in our study. Other methods of listening effort measurement, such as a dual-task paradigm (Xia et al., 2015) and functional near-infrared spectroscopy (fNIRS) (Andéol et al., 2017), also revealed spatial release from listening effort, although they used slightly different SNRs. Overall, all these studies have found a statistically significant negative relationship between SRM and LE, despite the differences in stimuli, target-masker configurations, type of masker, and type of measure used to assess listening effort.

Note, however, that Zekveld et al. (2014) and Thakkar et al. (2025) found no effect of spatial configuration on pupil responses in TH adults, indicating that although the availability of spatial hearing cues improved speech intelligibility, this benefit did not translate to release from listening effort. Instead, listening effort increased when SNR became more challenging, regardless of the spatial configuration. One reason that Zekveld et al. (2014) and Thakkar et al. (2025) did not observe release from effort with spatial separation could be that both studies have used the symmetrical masker configuration. Compared to our setup (both maskers on the same side), in which the monaural head shadow cues were available, in the symmetrical masking configuration monaural head shadow cues were absent. Without monaural head shadow cues, participants were required to rely solely on binaural cues. In our study, participants had access to both monaural and binaural cues from the spatial separation between the target and maskers, which may have been sufficient to yield a release from listening effort.

### 4.4 Listening Effort and Semantic Context Cues

One of the novelties of our study was that we manipulated the predictability of the target speech by using semantically coherent and anomalous sentences, which has not been done in any of the prior studies when examining the relationship between SRM and listening effort. We observed numerical differences in the peak pupil dilation between coherent and anomalous sentences, but these differences were not statistically significant. This comparison focused on semantic predictability while keeping spatial configuration constant. The absence of a significant effect may have been influenced by high variability between listeners, which may have reduced the power to detect group-level effects. The use of a single fixed +1 dB SNR may have contributed to this variability as well, as individual listeners are likely to experience different levels of task difficulty under the same SNR. In addition, although the behavioral data showed significant differences between coherent and anomalous sentences, the overall performance gaps were relatively modest, as compared to SRM, which may have limited the extent of observable pupil differences. Taken together, these findings point to two potential explanations for the absence of statistically significant group difference in peak dilation: inter-listener variability associated with the fixed SNR and the limited contrast between conditions, both of which are explored further below.

Previous studies that adopted individualized SNRs reported significant differences in peak pupil dilation between conditions (Koelewijn, Zekveld, Festen, Rönnberg, et al., 2012; Zekveld & Kramer, 2014). In these studies, individualized SNRs were derived from each listener’s speech reception threshold (SRT), which was measured by varying SNRs to determine the level yielding a target intelligibility. The target was typically defined as the SNR at which a listener could correctly repeat about 50 % of the speech material. By setting each participant’s SNR relative to their own SRT, those designs ensured a comparable level of task difficulty across individuals and reduced inter-listener variability. Under such individualized conditions, there was a statistically significant increase in peak pupil dilation at lower intelligibility levels, that is, at more challenging SNRs corresponding to each individual’s SRT. These results suggest that scaling task difficulty to individual performance levels can make pupil dilation a more sensitive marker of listening effort, potentially allowing the effects of semantic context to emerge.

Another factor influencing whether large group difference in peak pupil dilation occurs is the magnitude of contrast between listening conditions. Statistically significant changes in pupil dilation are typically observed only when the contrast between listening conditions is sufficiently large. For example, when SNR differs by several decibels (e.g., 0 dB vs. 6.5 dB), which leads to recognition scores ranging from about 60% to near ceiling. In contrast, smaller SNR steps (e.g. 0 dB vs. 3.5 dB) did not yield significant pupil differences (Zekveld & Kramer, 2014). In current study, behavioral performance differed modestly—about 20 % between coherent and anomalous sentences. These relatively small performance gaps suggest that the differences in task difficulty between conditions with varying semantic contexts might have been limited. This might explain why overall listening effort varied less with semantic predictability than intelligibility scores.

Although we did not observe statistically significant differences in peak pupil amplitude, analysis of the peak pupil latency revealed meaningful effects of semantic cues. In current study, peak pupil latency was significantly delayed for anomalous sentences compared to coherent sentences, indicating that listeners required additional time to sustain effort and retrieve target information when semantic context was absent. Although the mechanisms differ, reduced audibility in prior studies (Koelewijn, Zekveld, Festen, Rönnberg, et al., 2012; Zekveld & Kramer, 2014) versus reduced predictability here, the consistent finding of delayed peak latency indicates that listeners reach peak pupil dilation later under more challenging listening conditions, such as lower SNRs or reduced semantic predictability, reflecting sustained cognitive effort during these difficult tasks. Longer latencies therefore, appear to be a robust marker of prolonged processing effort, observable across both acoustically and linguistically demanding tasks. This effect extends across modalities, including EEG, where latency frequently serves as a more reliable index of cognitive demand than amplitude measures (Borjigin et al., 2022).

Most earlier studies examined only one type of sentence material, either coherent (semantically predictable) or anomalous (semantically unpredictable), without directly comparing the influence of contextual cues on listening effort (Koelewijn, Zekveld, Festen, Rönnberg, et al., 2012; Zekveld & Kramer, 2014). Current study extended previous work by contrasting two sentence types while holding acoustic difficulty constant. Our manipulation of contextual information changed the linguistic predictability of the sentences. By varying predictability rather than acoustic difficulty, the task places a cognitive demand on how efficiently listeners can use context to guide comprehension, which helps them process the acoustic signal more efficiently and with less effort. In other words, we compared listening behavior when sentences were meaningful and predictable (coherent) versus when they were grammatically correct but lacked contextual cues (anomalous).

Through this design, our study captures how differences in semantic context influenced not only the amount of listening effort but also how long that listening effort was sustained. These two aspects were reflected in distinct pupil measures: peak pupil dilation indicates the magnitude of effort, whereas latency reflects how long that effort was sustained. In the current study, peak amplitude did not differ statistically significantly between coherent and anomalous sentences, suggesting that removing contextual information might not have increased the overall intensity of effort. However, we observed a significant difference in latency: listeners sustained effort for a longer period when semantic cues were unavailable. These findings show that semantic predictability influences the timing and persistence of listening effort: when predictability is low, listeners maintain effort for longer periods to achieve comprehension, even though the overall level of effort might be similar.

### 4.5 Limitations and Future Directions

In the current study, we found that listening effort can further explain individual differences in speech recognition with varying levels of spatial separation and semantic cues in TH adults. However, because we evaluated listening effort and SRM at only one SNR,, one immediate future direction is to examine the interaction of spatial separation and semantic cues, LE, and SNR, in a wide range of intelligibility levels. This would reveal how cognitive processing load associated with listening effort scales as task demand shift. Another future direction is to test TH adults with cochlear implant simulation and adults who are fitted with hearing aids or cochlear implants. This extension is important because access to spatial hearing cues is often degraded in these populations. Testing these groups will determine whether the interactions between spatial cues and sentence-level semantic context observed in normal-hearing listeners generalize to individuals with degraded auditory input. Ultimately, this work will clarify whether restoring or enhancing spatial cues provides measurable clinical benefits—specifically, whether it reduces listening effort in addition to improving speech intelligibility. This extension is important because access to spatial hearing cues is often degraded in these populations, and it remains unclear whether restoring or enhancing spatial cues would reduce listening effort in addition to improving speech intelligibility. Testing these groups would help determine whether the interaction between spatial cues and sentence-level semantic context observed in normal-hearing listeners generalizes to individuals with degraded auditory input, and whether spatial hearing restoration provides measurable benefits for both speech understanding and listening effort.

## Statements and Declarations

### Ethical Considerations

All experimental procedures were conducted in accordance with National Institutes of Health regulations and approved by the Health Sciences Institutional Review Board (#2016-0226) at the University of Wisconsin– Madison.

### Consent to Participate

All participants reviewed and signed the consent form before participating in the study.

### Consent for Publication

Not applicable.

### Data Availability

The data that support the findings of this study will be made available on Open Science Framework.

